# BTB-TAZ domain protein MdBT2 modulates malate accumulation by targeting a bHLH transcription factor for degradation in response to nitrate

**DOI:** 10.1101/813832

**Authors:** Quan-Yan Zhang, Kai-Di Gu, Lailiang Cheng, Jia-Hui Wang, Jian-Qiang Yu, Xiao-Fei Wang, Chun-Xiang You, Da-Gang Hu, Yu-Jin Hao

## Abstract

Excessive application of nitrate, an essential macronutrient and a signal regulating diverse physiological processes, decreases malate accumulation in apple fruit, but the underlying mechanism remains poorly understood. Here, we show that an apple BTB/TAZ protein MdBT2 is involved in regulating malate accumulation and vacuolar pH in response to nitrate. *In vitro* and *in vivo* assays indicate that MdBT2 interacts directly with and ubiquitinates a bHLH transcription factor, MdCIbHLH1, via the ubiquitin/26S proteasome pathway in response to nitrate. This ubiquitination results in the degradation of MdCIbHLH1 protein and reduces the transcription of MdCIbHLH1-targeted genes involved in malate accumulation and vacuolar acidification including *MdVHA-A* encoding a vacuolar H^+^-ATPase gene, and *MdVHP1* encoding a vacuolar H^+^-pyrophosphatase gene, as well as *MdALMT9* encoding a aluminum-activated malate transporter gene. A series of transgenic analyses in apple materials including fruits, plantlets and calli demonstrate that MdBT2 controls nitrate-mediated malate accumulation and vacuolar pH at least partially, if not completely, via regulating the MdCIbHLH1 protein level. Taken together, these findings reveal that MdBT2 regulates the stability of MdCIbHLH1 via ubiquitination in response to nitrate, which in succession transcriptionally reduces the expression of malate-associated genes, thereby controlling malate accumulation and vacuolar acidification in apples under high nitrate supply.

## Introduction

Malate, as a key metabolite, plays a vital role in plant metabolism, pH homeostasis, nutrient uptake, osmotic adjustment and abiotic stress resistance (Fernie and Martinoia, 2009; Finkemeier and Sweetlove, 2009; Bai et al., 2015; Hu et al., 2017). Cellular malate accumulation also largely determines the acidity and perception of sweetness of fleshy fruits and their processed products (Yao et al., 2007; Ye et al., 2017; Butelli et al., 2019; Ma et al., 2019). The majority of malate in the parenchyma cells of fleshy fruits is in the vacuole (Yamaki, 1984). Malate accumulation is a complex process involving synthesis, degradation and transport of malate. Although malate metabolism can alter fruit malate level (Sweetman et al., 2009; Centeno et al., 2011), it appears that transport of malate from the cytosol into the vacuole is the step that largely controls malate accumulation (Etienne et al., 2013; Hu et al., 2016a; Ma et al., 2019).

Vacuolar proton pumps and malate transporters are crucial for malate transport into the vacuole. Vacuolar proton pumps, such as V-ATPase and V-PPase, acidify the vacuoles by pumping protons across the tonoplast. The resulting low pH protonates any malate that crosses the tonoplast from the cytosol, effectively trapping malate in the acid form. This “acid trap” mechanism maintains the malate concentration gradient across the tonoplast for its facilitated diffusion (Martinoia et al., 2007; Etienne et al., 2013; Eisenach et al., 2014; Hu et al., 2016a). Aluminum-activated malate transporter/channel (ALMT) as well as tonoplast dicarboxylate transporter (tDT) enables the transport of malate across the tonoplast (Kovermann et al., 2007; Ma et al., 2015; Ye et al., 2017). *ALMT9* underlies a major malic acid locus, *Ma* in apple (Bai et al., 2012; Khan et al., 2013) and a major QTL for malate level in tomato (Jie et al., 2017).

Upstream of these proton pumps and transporters are various transcription factors (TFs) that regulate their expression. For example, SlWRKY42 directly binds and affects malate transporter SlALMT9 to negatively regulate malate content in tomato fruit (Ye et al., 2017). Furthermore, the MYB TFs together with bHLH TFs and WD40 proteins form MBW complexes to transcriptionally activate the expression of vacuolar proton pumps and malate transporter genes, thereby promoting malate accumulation in the vacuole (Xie et al., 2012; Hu et al., 2016a, 2017). In apple, MdMYB73 binds directly to the promoters of *MdVHA-A*, *MdVHP1* and *MdALMT9* to regulate malate accumulation (Hu et al., 2017). MdCIbHLH1, a bHLH transcription factor homologous to *Arabidopsis* ICE1, interacts with MdMYB73 and enhances its effects on downstream target genes to regulate malate accumulation in apple (Feng et al., 2012; Hu et al., 2017).

Malate accumulation in fleshy fruits is also affected by environmental factors and management practices such as nutrient availability, salinity, temperature, and irrigation (Wu et al., 2002; Burdon et al., 2007; Thakur and Singh, 2012; Etienne et al., 2013). In response to environmental stresses such as low temperature and salinity, plants accumulate malate as a metabolite in coping with these stresses, consequently altering fruit acidity level (Ruffner et al., 1976; Friemert et al., 1988; Hu et al., 2016b). Nitrate is an essential macronutrient for plant growth and development, and also serves as a signal regulating many physiological processes (Liu et al., 2017; Maeda et al., 2018). High nitrate supply has been reported to reduce malate accumulation (Spironello et al., 2004), but how nitrate affects the malate accumulation remains poorly understood.

The BTB-TAZ proteins respond to diverse environmental stimuli such as nutrients and stress and internal signals such as hormones in plants (Mandadi et al., 2009). We recently found that BT2 as a BTB-TAZ protein responds to nitrate in modulating anthocyanin synthesis and accumulation in apple (Wang et al., 2018). In this work, we showed that MdBT2 protein regulates malate accumulation in response to nitrate. A yeast two-hybrid screening library allowed for identification of MdCIbHLH1 as a candidate protein for MdBT2 interaction. Subsequently, we uncovered that MdBT2 regulates the stability of MdCIbHLH1 via ubiquitination in response to nitrate, which in succession transcriptionally reduces the expression of several malate-associated genes in altering malate accumulation in apple.

## Results

### Exogenous application of nitrate reduces malate accumulation

To determine if nitrate has any effect on malate accumulation, we provided apple calli and plantlets with different nitrate concentrations (0 to 5 mM) and measured malate contents in various tissues first. High concentrations of nitrate were found to significantly reduce tissue malate accumulation (Supplemental Fig. S1).

To confirm the effect of nitrate on malate accumulation in fruits, we treated ‘Royal Gala’ apple fruits with exogenous nitrate from 0 to 5 mM for further analysis (Fig. 1A). Nitrate application led to a significant reduction in fruit malate level (Fig. 1C). The expression levels of malate-related genes, including vacuolar proton pumps *MdVHA-A*, *MdVHA-Bs* and *MdVHP1*, and the vacuolar malate channel *MdALMT9* as well as malate-associated MYB TF *MdMYB73*, were significantly decreased when nitrate concentration increased from 0 to 5 mM (Fig. 1B). Subsequently, we measured the hydrolytic and proton-pumping activities of V-ATPase and V-PPase activity. V-ATPase hydrolytic activity and proton-pumping activity and V-PPase activity were decreased to 1/2, 2/3, and 1/3 of the 0 mM N control, respectively, in the 5 mM N treatment (Fig. 1, D-F). Furthermore, measurements of vacuolar pH with the ratiometric fluorescent pH indicator 2’,7’-bis-(2-carboxyethyl)-5-(6)-carboxyfluorescein (BCECF) showed that the average vacuolar pH was increased from 3.6 in 0 mM N to 3.7 and 4.1, respectively, in the 0.5 mM and 5 mM N treatments (Fig. 1, G-H). These results suggest that high nitrate level reduces vacuolar acidification by decreasing the proton-pumping activities of both V-ATPase and V-PPase in apple.

**Figure 1.**
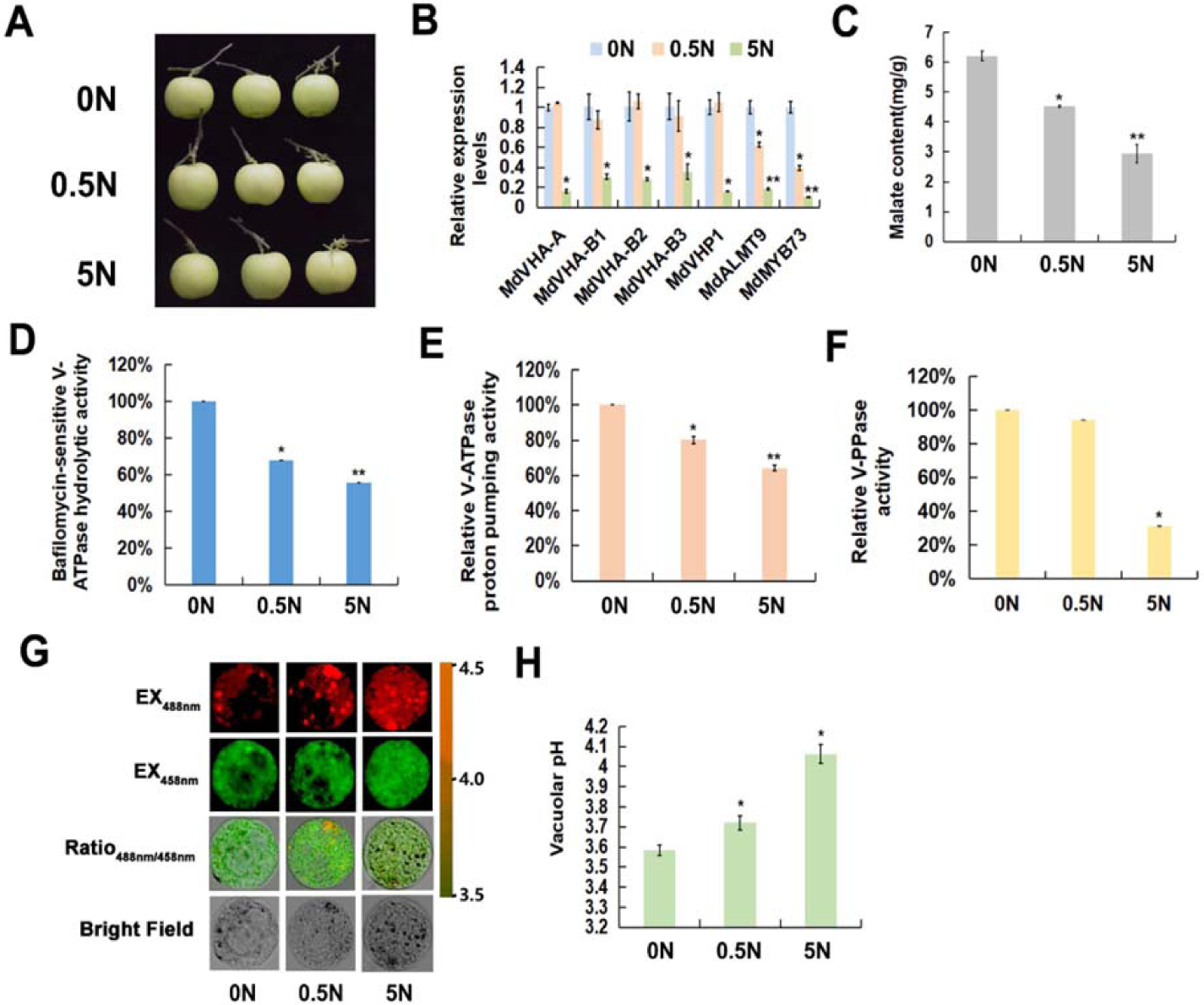
Exogenous application of nitrate reduces malate accumulation. **A,** The ‘*Royal Gala*’ apple fruits were treated with different concentrations of exogenous nitrate (including 0 mM, 0.5 mM and 5 mM nitrate). **B,** The expression of malate-related genes under different concentrations of exogenous nitrate using qPCR. The genes included about vacuolar proton pumps *MdVHA-A*, *MdVHA-Bs* and *MdVHP1*, and the vacuolar malate channel *MdALMT9* as well as malate-associated MYB TF *MdMYB73*. **C,** Malate contents under different concentrations of exogenous nitrate. **D,** Bafilomycin A1-sensitive ATP hydrolytic activity of V-ATPase under different concentrations of exogenous nitrate. **E,** Proton-pumping activity of V-ATPase under different concentrations of exogenous nitrate was measured by tracking the ATP-dependent quenching of acridine orange using a fluorescence spectrometer with excitation at 493 nm and emission at 545 nm. **F,** V-PPase activity under different concentrations of exogenous nitrate. **G,** Emission intensities of protoplast vacuoles under different concentrations of exogenous nitrate loaded with BCECF at 488 nm (first column) and 458 nm (second column). Scale bar = 10 um. **H,** Quantification of the pH in vacuoles under different concentrations of exogenous nitrate. Significant difference was detected by *t*-test. *P < 0.05, **P < 0.01.

### BTB/TAZ protein MdBT2 controls malate accumulation in response to nitrate

Considering that BT2 as a BTB-TAZ protein responds to nitrate in plants (Mandadi et al., 2009; Wang et al., 2018), we explored the potential role of BT2 in nitrate-modulated malate accumulation. We first looked at the expression of *MdBT2* in response to nitrate and found that nitrate application increased its transcript level (Supplemental Fig. S2). We then generated *35S::anti-MdBT2* RNAi transgenic apple calli (35S::anti-MdBT2) (Supplemental Fig. S2), and detected higher malate levels in the anti-*MdBT2* calli.

To confirm the role of MdBT2 in malate accumulation in response to nitrate, transgenic apple plantlets overexpressing *MdBT2* (MdBT2-OX) were treated with 0 mM N (0 N-MdBT2-OX), while *antiMdBT2* transgenic apple plantlets (antiMdBT2) were treated with 5 mM N (5N-antiMdBT2), along with wild-type (WT) control under 0 mM or 5 mM nitrate. 5 mM nitrate decreased malate accumulation in WT plantlets; MdBT2-OX reduced the malate level at 0 mM nitrate whereas antiMdBT2 increased the malate level at 5 mM nitrate (Fig. 2A).

**Figure 2.**
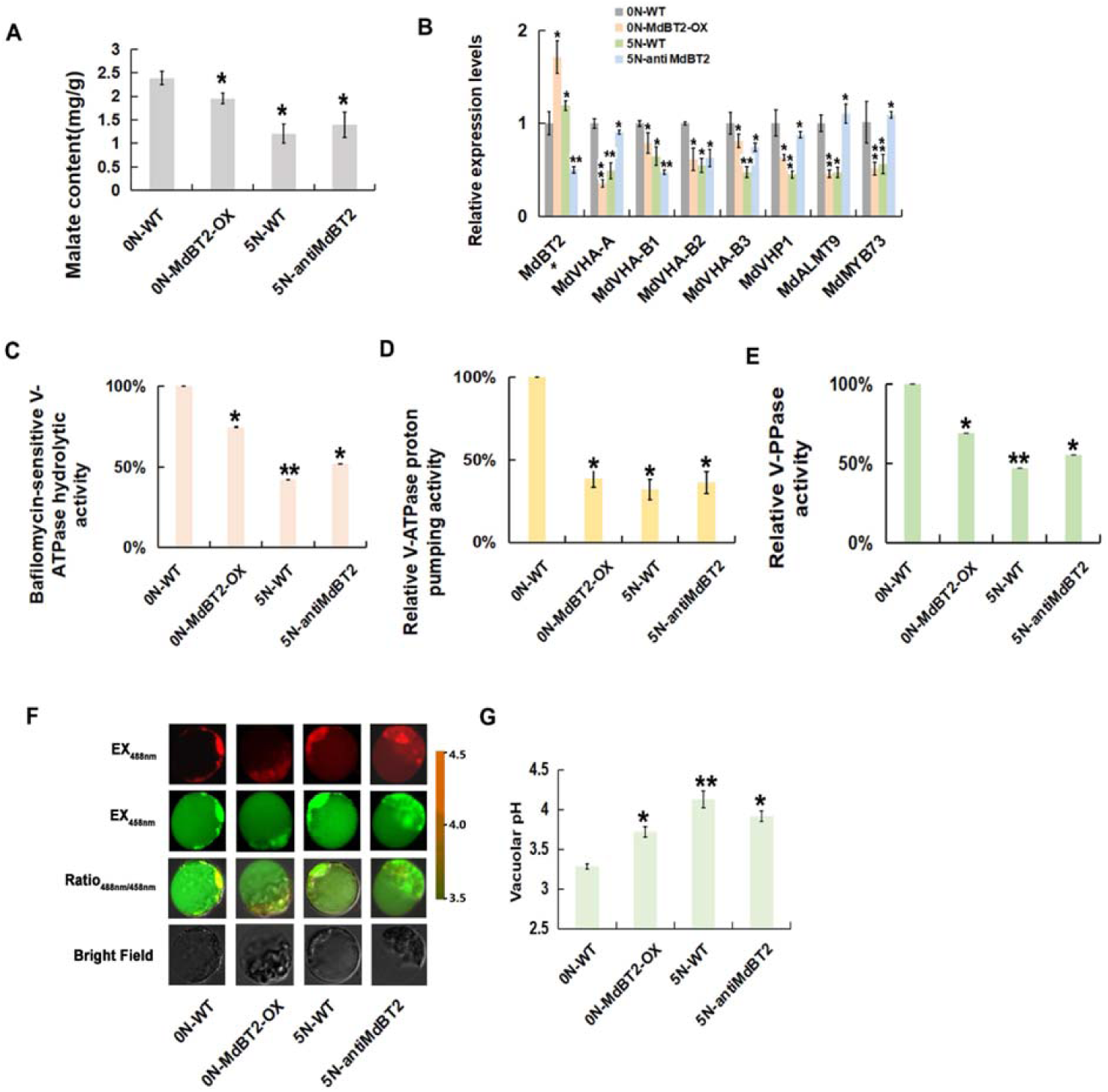
BTB/TAZ protein MdBT2 controls malate accumulation in response to nitrate. **A,** Malate contents in WT and transgenic apple plantlets under different concentrations of exogenous nitrate. ‘GL-3’ plantlets were treated with 0 N (0 N-WT). MdBT2 overexpression transgenic apple plantlets were treated with 0 N (0 N-MdBT2-OX). The antiMdBT2 transgenic apple plantlets were treated with 5 N (5 N-antiMdBT2). **B,** Expression of *MdVHA-A*, *MdVHP1* and *MdALMT9* in WT and transgenic apple plantlets under different concentrations of exogenous nitrate using qPCR. **C to E,** The hydrolytic and proton-pumping activities of V-ATPase and V-PPase in WT and transgenic apple plantlets under different concentrations of exogenous nitrate. **F,** Emission intensities of protoplast vacuoles loaded with BCECF at 488 nm and 458nm. Scale bar = 10 um. **G,** Quantification of the pH in vacuoles. Significant difference was detected by *t*-test. *P < 0.05, **P < 0.01.

Correspondingly, 5 mM nitrate inhibited the expression of malate-associated genes including *MdVHA-A*, *MdVHA-Bs*, *MdVHP1* and *MdALMT9*, and V-ATPase hydrolytic and proton-pumping activities as well as V-PPase activity (Fig. 2, B-E); overexpression of *MdBT2* decreased the expression of these genes and activities of related enzymes and increased vacuolar pH at 0 mM nitrate whereas antisense repression of *MdBT2* increased the expression of the genes and activities of related enzymes and decreased vacuolar pH at 5 mM nitrate (Fig. 2, B-G).

Taken together, these results demonstrate that BTB/TAZ protein MdBT2 modulates malate accumulation in response to nitrate in apple.

### MdBT2 physically interacts with MdCIbHLH1 via the conserved BTB/BACK domain

To explore the regulatory mechanism by which MdBT2 is involved in malate accumulation in apple, possible MdBT2-interacting proteins were screened using MdBT2 as a bait through a yeast two hybrid (Y2H) cDNA library. MdCIbHLH1, a basic helix-loop-helix (bHLH) TF that was previously identified to be involved in malate accumulation and cold resistance, was chosen as a candidate. To determine whether MdBT2 interacts with MdCIbHLH1 protein, Y2H assays were performed. As MdBT2 protein contains three conserved domains, BTB, BACK and ZnF_TAZ, we divided the full-length cDNA of *MdBT2* gene into four fragments, i.e., MdBT2^BTB/BACK^, and MdBT2^BTB^, as well as MdBT2^BACK^ and MdBT2^BACK/ZnF_TAZ^. Subsequently, the full length cDNA and four truncated mutants of *MdBT2* gene were inserted into the pGBT9 vector, independently, as the bait vectors. Meanwhile, the full-length MdCIbHLH1 cDNA was inserted into the pGAD424 as the prey vector. The different combinations of bait and prey vectors were transformed into yeast for Y2H assays. The full-length MdBT2 interacted with the full-length MdCIbHLH1 protein (Fig. 3A). Moreover, MdCIbHLH1 interacted with the truncated mutant MdBT2^BTB/BACK^ but not with others (Fig. 3A). These results suggest that MdBT2 *in vitro* interacts with MdCIbHLH1 via the conserved BTB/BACK domain. In addition, a GST pull-down assay showed that a GST-tagged MdBT2 physically interacted with a His-tagged MdCIbHLH1 *in vitro* (Fig. 3B).

**Figure 3.**
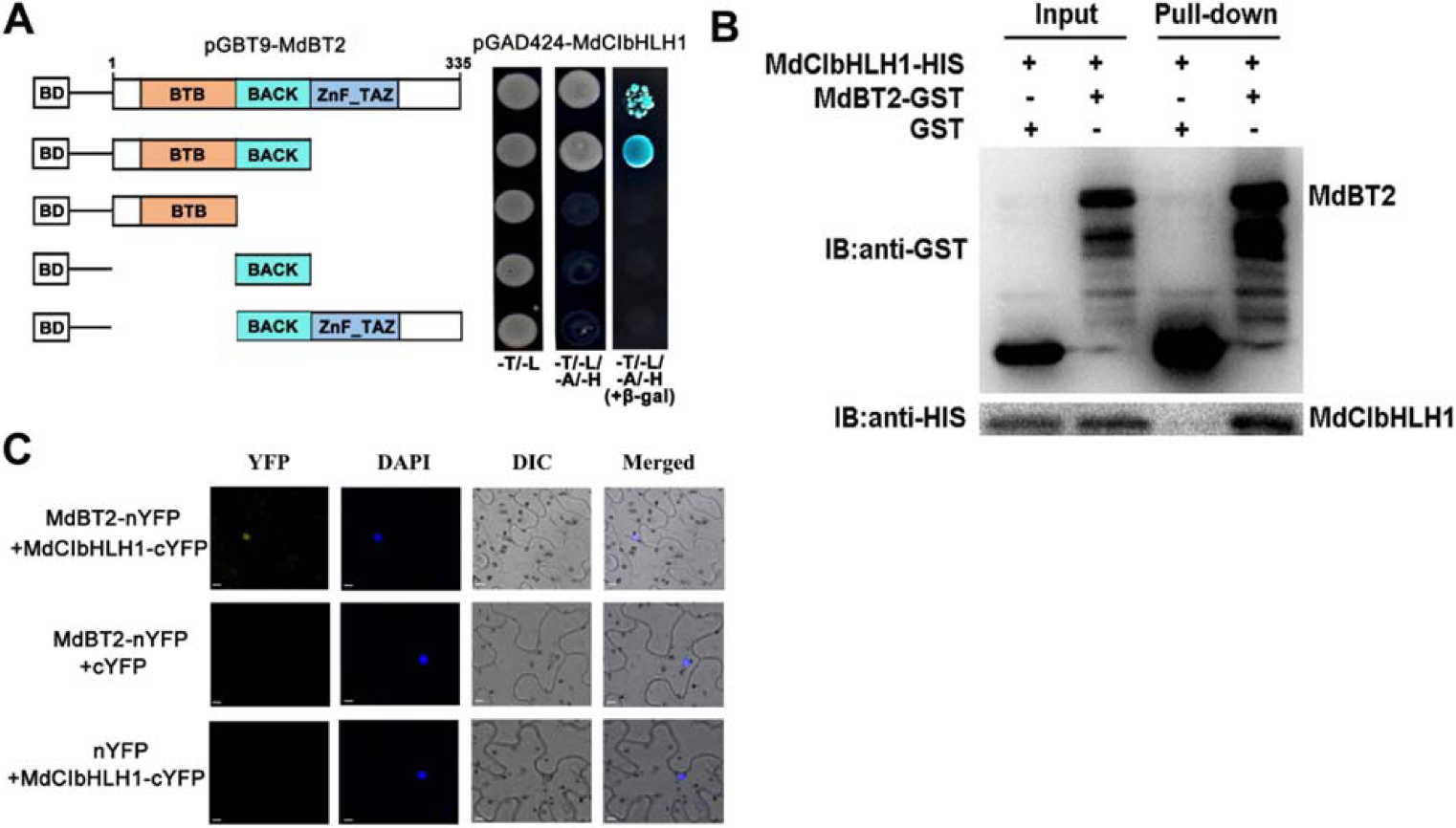
MdBT2 physically interacts with MdCIbHLH1 via the conserved BTB/BACK domain. **A,** MdBT2 interacted with MdCIbHLH1 in a Y2H assay. The full-length and the BTB/BACK domain of MdBT2 interacted with the MdCIbHLH1 protein. **B,** Pull-down assays of the interaction between MdBT2 and MdCIbHLH1. MdBT2-GST was expressed by *E. coli* resulted in the precipitation of MdCIbHLH1-HIS using anti-GST antibody and anti-HIS antibody, respectively. GST alone was used as the control. **C,** BiFC assay showed the interaction between MdBT2 and MdCIbHLH1 using tobacco leaf cells. MdBT2-nYFP and MdCIbHLH1-cYFP interacted in the nucleus of tobacco leaf cells. Scale bar = 10 um.

To further confirm the interaction between MdBT2 and MdCIbHLH1, an *in vivo* Bimolecular fluorescence complementation (BiFC) assay using tobacco leaf cells was conducted. MdBT2 and MdCIbHLH1 were ligated to the N-terminal and C-terminal of YFP, respectively, to generate the constructs MdBT2-nYFP and MdCIbHLH1-cYFP. Subsequently, these constructs including empty vectors with pairwise combinations were transiently transformed into tobacco leaf cells by *Agrobacterium*-mediated method. A strong YFP signal was observed in the nucleus of tobacco leaf cells in the combination of MdBT2-nYFP and MdCIbHLH1-cYFP constructs, with no signal detected in the other two combinations (MdBT2-nYFP+cYFP and nYFP+MdCIbHLH1-cYFP) as negative controls (Fig. 3C). These results indicate that MdBT2 physically interacts with the MdCIbHLH1 protein.

### MdCIbHLH1 is involved in regulating malate accumulation

It is known that MdCIbHLH1 interacts with MdMYB73 to modulate malate accumulation by directly activating vacuolar transporter genes including *MdALMT9*, *MdVHA-A* and *MdVHP1* in apple (Hu et al., 2017). To further verify the functions of MdCIbHLH1 in the regulation of malate accumulation in apple fruits, the *35S::MdCIbHLH1* transgenic apple plantlets (MdCIbHLH1-OXs) obtained by Feng et al. (2012) were grafted onto rootstock M9T337, and transgenic apple fruits were harvested at different stages during fruit development in the third growing season. Determination of malate contents revealed that fruits of the three *MdCIbHLH1* transgenic lines (MdCIbHLH1-OVXs) had much higher malate levels than the wild-type control (*‘GALA’*) throughout fruit development, particularly at 40 days after flowering (DAF) (Fig. 4A). Subsequently, 40 DAF apple fruits were used for further analysis (Fig. 4B).

**Figure 4.**
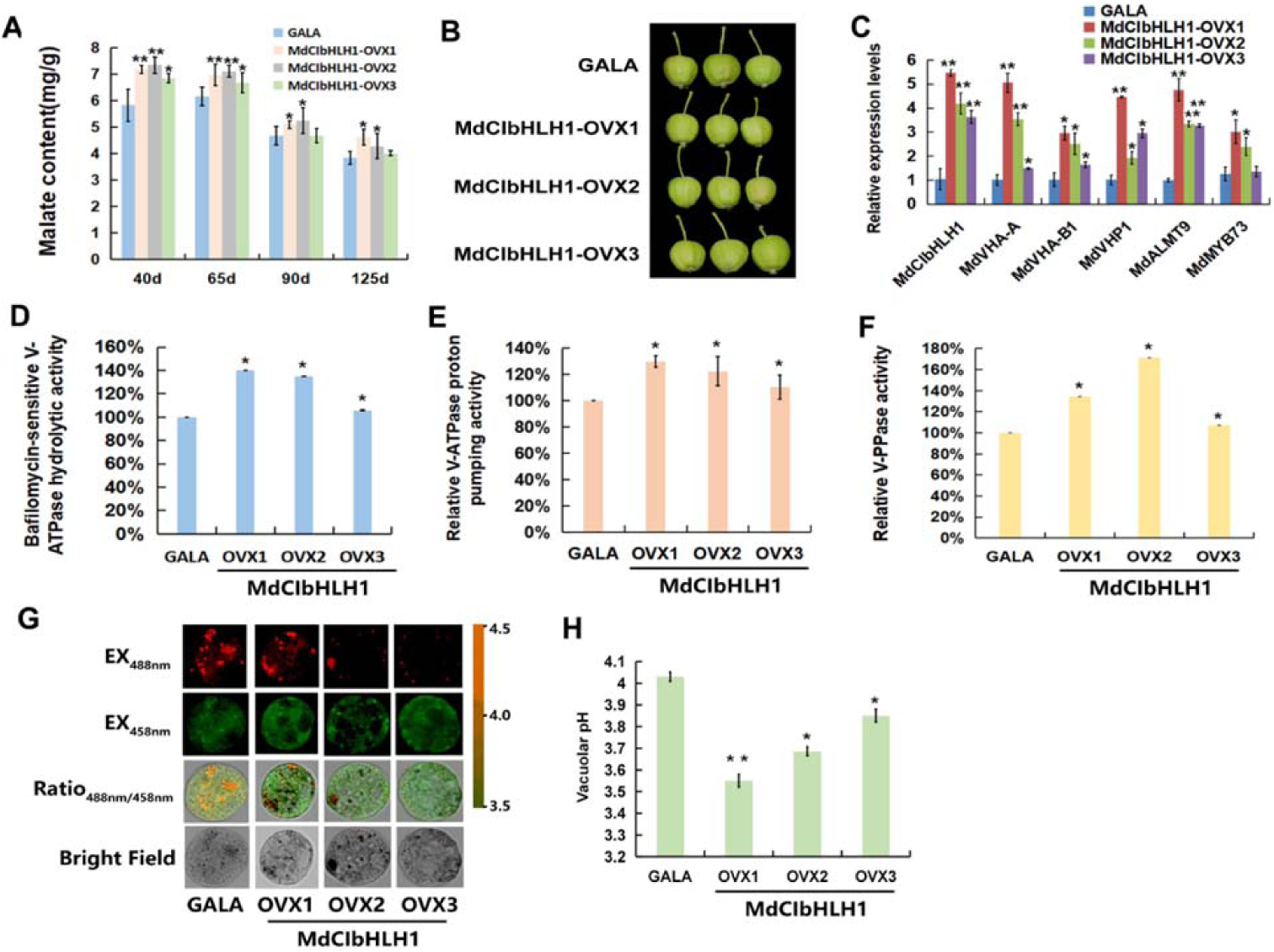
MdCIbHLH1 is involved in regulating of malate accumulation. **A,** Malate contents in ‘*GALA*’ and *MdCIbHLH1-OVXs* transgenic apple fruits (MdCIbHLH1-OVXs) in the whole fruit developmental stages. **B,** The ‘*GALA*’ and MdCIbHLH1-OVXs taken in 40 DAF. **C,** The expression of *MdVHA-A*, *MdVHA-B1*, *MdVHP1*, *MdALMT9* and *MdMYB73* in *‘GALA’* and MdCIbHLH1-OVXs. **D to F,** The hydrolytic and proton-pumping activities of V-ATPase and V-PPase in ‘*GALA*’ and MdCIbHLH1-OVXs. **G,** Emission intensities of protoplast vacuoles in ‘*GALA*’ and MdCIbHLH1-OVXs with BCECF at 488 nm and 458 nm. Scale bar = 10 um. **H,** Quantification of the pH in vacuoles in ‘*GALA*’ and MdCIbHLH1-OVXs. Significant difference was detected by *t*-test. *P < 0.05, **P < 0.01.

Transcript levels of *MdVHA-A*, *MdVHA-B1* and *MdVHP1*, as well as *MdALMT9* and *MdMYB73*, were significantly higher in the three independent MdCIbHLH1-OVXs than in *‘GALA’* fruits control (Fig. 4C). V-ATPase and V-PPase activities were 1.2- to 1.8-fold greater in MdCIbHLH1-OVXs than in ‘*GALA*’ fruits (Fig. 4, D-F). Furthermore, we used BCECF to illustrate the effects of V-ATPase and V-PPase activities on vacuolar pH. As shown in Fig. 4, G-H, the average vacuolar pH in ‘*GALA*’ fruits was 4.03 while those of the three MdCIbHLH1-OVXs were 3.56, 3.68 and 3.84, respectively. Hence, higher vacuolar H^+^ concentrations in *MdCIbHLH1*-expressing fruits, indicated by the lower pH values, further supports the conclusion that MdCIbHLH1 regulates malate accumulation by altering the tonoplast V-ATPase and V-PPase activities.

### MdBT2 affects the stability of MdCIbHLH1 protein

Earlier work showed that ubiquitination-related MdBT2 scaffold protein targets MYB and bHLH TFs for degradation, thereby regulating plant senescence and anthocyanin biosynthesis (Wang et al., 2018; An et al., 2019). Considering the interaction between MdBT2 and MdCIbHLH1, we predicted that MdBT2 probably affects the stability of MdCIbHLH1 protein. To verify this, cell-free degradation assays of the prokaryon-expressed and purified MdCIbHLH1-HIS fusion proteins were conducted using protein samples extracted from wild-type (WT), 35S::MdBT2 transgenic apple calli (35S::MdBT2) and 35S::anti-MdBT2. The MdCIbHLH1-HIS protein was more rapidly degraded in the protein extract of the 35S::MdBT2 than in that of the WT (Figure 5A), whereas the protein was more stable in the protein extract of 35S::anti-MdBT2 compared to the WT (Fig. 5A). These results suggest that MdBT2 promotes the degradation of the MdCIbHLH1 protein.

**Figure 5.**
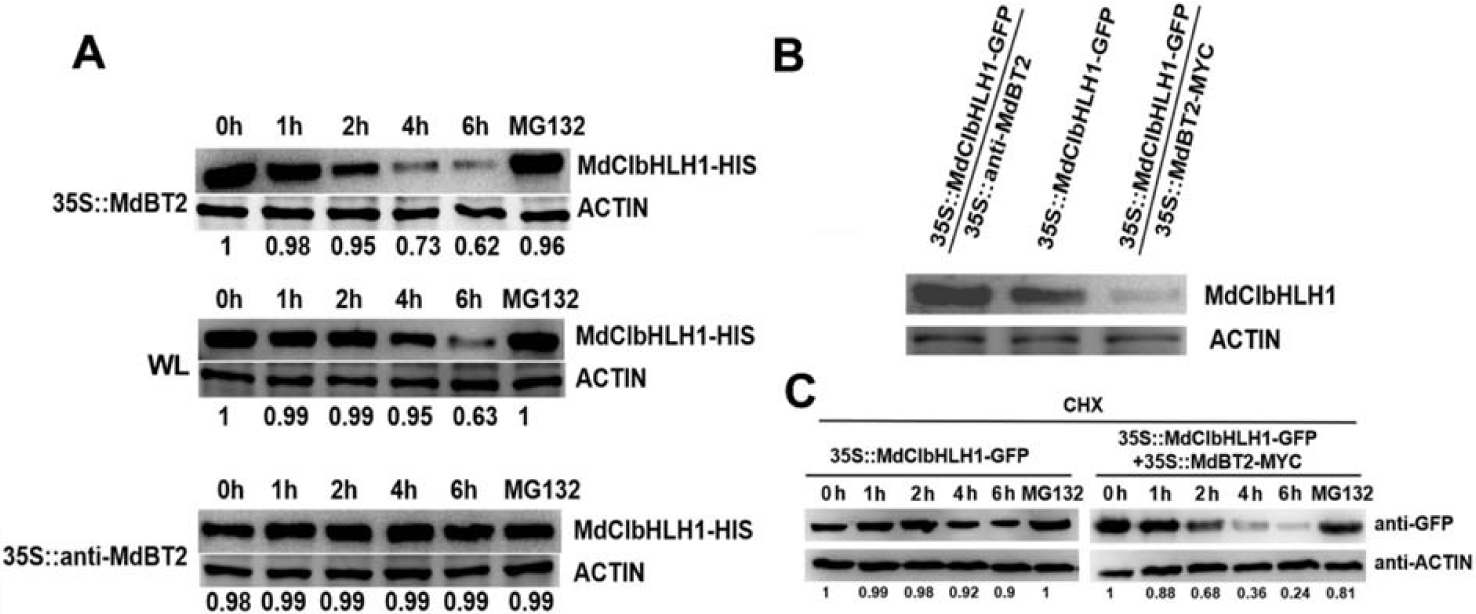
MdBT2 affects the stability of MdCIbHLH1 protein. **A,** The cell-free degradation assays of the prokaryon-expressed and purified MdCIbHLH1-HIS fusion proteins were using protein samples extracted from WL, 35S::MdBT2 and 35S::anti-MdBT2. **B,** Three types of transgenic calli, that is, 35S::MdCIbHLH1-GFP, 35S::MdBT2-MYC + 35S::MdCIbHLH1-GFP and 35S::anti-MdBT2 + 35S::MdCIbHLH1-GFP were obtained and used for immunoblotting assays with an anti-GFP antibody. **C,** The cell-free degradation assays were also performed using 35S::MdCIbHLH1-GFP and 35S::MdBT2-MYC + 35S::MdCIbHLH1-GFP. Total proteins were analyzed by immunoblotting using the anti-GFP antibody. ACTIN was used as the loading control.

To further examine whether MdBT2 influences the degradation of the MdCIbHLH1 protein *in vivo*, three types of transgenic calli, 35S::MdCIbHLH1-GFP, 35S::MdBT2-MYC + 35S::MdCIbHLH1-GFP, and 35S::anti-MdBT2 + 35S::MdCIbHLH1-GFP were obtained and used for immunoblotting assays with an anti-GFP antibody. The abundance of MdCIbHLH1 was lower in 35S::MdBT2-MYC + 35S::MdCIbHLH1-GFP, but higher in 35S::anti-MdBT2 + 35S::MdCIbHLH1-GFP than in the WT (Fig. 5B). Meanwhile, cell-free degradation assays were also performed using 35S::MdCIbHLH1-GFP and 35S::MdBT2-MYC + 35S::MdCIbHLH1-GFP. As shown in Fig. 5C, the MdCIbHLH1-GFP protein in 35S::MdBT2-MYC + 35S::MdCIbHLH1-GFP was degraded much greatly than in 35S::MdCIbHLH1-GFP, just as they did *in vitro*.

### MdBT2 is involved in the ubiquitination and degradation of MdCIbHLH1 proteins

To examine whether MdCIbHLH1 is degraded through a 26S proteasomal pathway, 35S::MdCIbHLH1-GFP treated with the translational inhibitor cycloheximide (CHX) and the proteasomal inhibitor MG132 were used for immunoblotting assays with an anti-GFP antibody. The results show that MdCIbHLH1 protein accumulation was almost completely inhibited by CHX in the 35S::MdCIbHLH1-GFP (Fig. 6A). By contrast, MdCIbHLH1 protein highly accumulated in the transgenic calli treated with MG132 (Fig. 6A), suggesting that the MdCIbHLH1 protein is degraded in a 26S proteasome-dependent manner.

**Figure 6.**
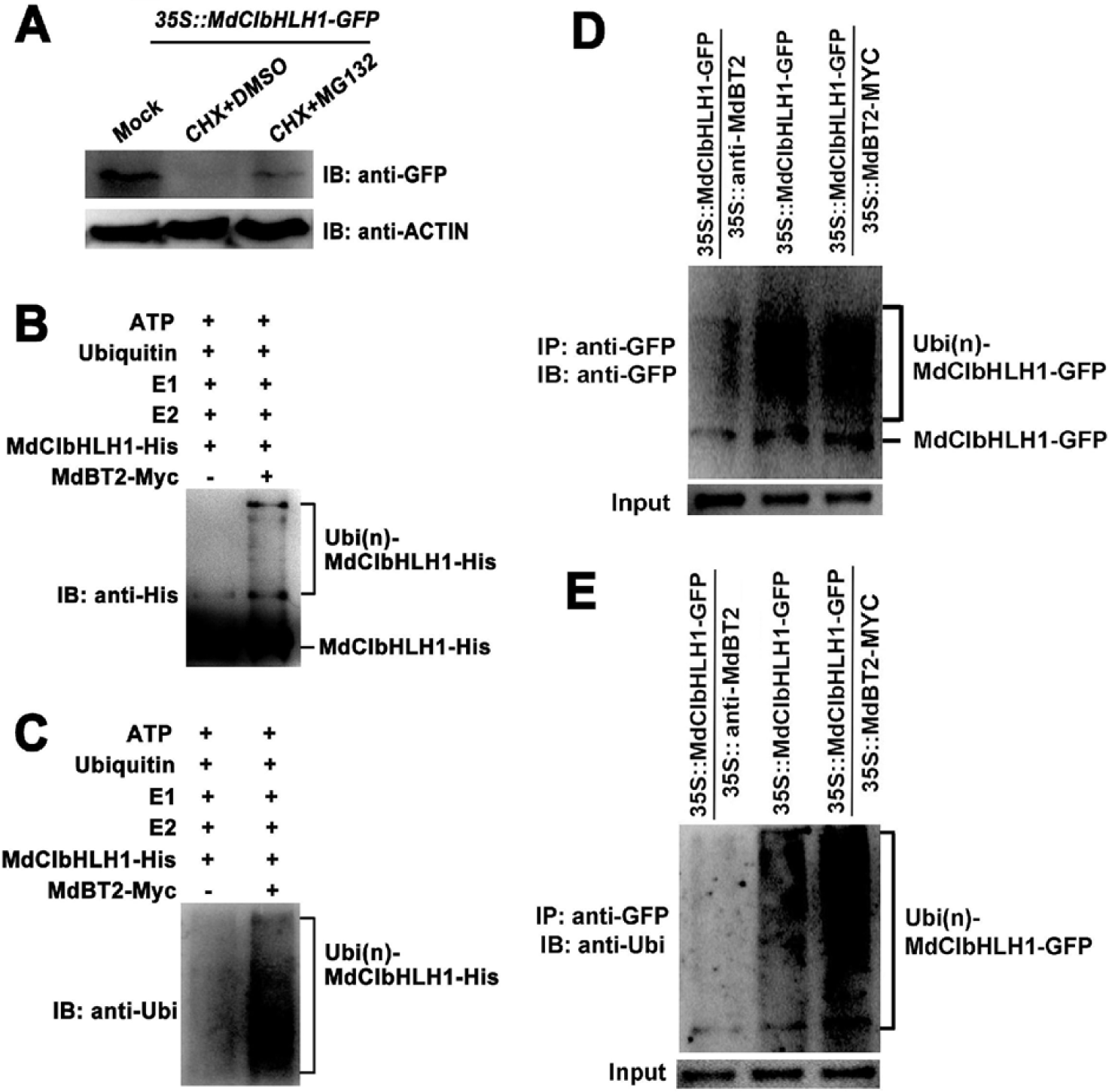
MdBT2 is involved in the ubiquitination and degradation of MdCIbHLH1 proteins. **A,** 35S::MdCIbHLH1-GFP were treated with CHX and MG132 for 6h and then measured the MdCIbHLH1-GFP protein level using anti-GFP antibody. **B to E,** MdBT2 promoted the ubiquitination of MdCIbHLH1 protein *in vitro* and *vivo*. **B and C,** The MdBT2-MYC active protein extracted from 35S::MdBT2-MYC were co-incubated with prokaryon-expressed and purified MdCIbHLH1-HIS protein, E1, E2 and ubi *in vitro* for immunoprecipitation using anti-HIS and anti-ubi antibodies. **D and E,** Ubiquitination assays *in vivo* were measured using 35S::MdCIbHLH1-GFP, 35S::MdCIbHLH1-GFP + 35S::MdBT2-MYC and 35S::MdCIbHLH1-GFP + 35S::anti-MdBT2. The anti-GFP antibody and anti-ubi antibodies were used to detect the MdCIbHLH1-GFP proteins accumulation. IP, Immunoprecipitated. IB, immunoblotted.

26S proteasome-mediated protein degradation is generally associated with ubiquitination modifications. To examine if MdBT2 participates in the ubiquitination of the MdCIbHLH1 protein, the MdBT2-MYC active protein extracted from 35S::MdBT2-MYC were co-incubated with prokaryon-expressed and purified MdCIbHLH1-HIS protein, E1, E2 and ubi *in vitro* for immunoprecipitation using anti-HIS and anti-ubi antibodies. A higher amount of high-molecular mass forms of MdCIbHLH1, that is, polyubiquitinated MdCIbHLH1 (Ubi(n)-MdCIbHLH1–HIS), was detected when supplemented with MdBT2-MYC active protein (Fig. 6, B and C). Therefore, it seems that MdBT2 is essential for the ubiquitination of MdCIbHLH1 proteins.

Subsequently, *in vivo* ubiquitination assays were conducted using 35S::MdCIbHLH1-GFP, 35S::MdCIbHLH1-GFP + 35S::MdBT2-MYC and 35S::MdCIbHLH1-GFP + 35S::anti-MdBT2 (Fig. 6, D and E). The anti-GFP antibody and anti-ubi antibodies were used to detect the MdCIbHLH1-GFP protein accumulation. The abundance of polyubiquitinated MdCIbHLH1-GFP was greater in 35S::MdCIbHLH1-GFP + 35S::MdBT2-MYC than in 35S::MdCIbHLH1-GFP, but lower in 35S::MdCIbHLH1-GFP + 35S::anti-MdBT2 than in 35S::MdCIbHLH1-GFP (Fig. 6, D and E). These results suggest that the ubiquitination and degradation of the MdCIbHLH1 protein was mediated by MdBT2.

### Nitrate participates in the regulation of MdBT2-mediated MdCIbHLH1 protein stability

To determine whether nitrate is involved in the regulation of MdBT2-mediated MdCIbHLH1 protein stability, the 35S::MdCIbHLH1-GFP calli were provided with nitrate (+N) or without nitrate (-N). The total proteins extracted from these calli samples were then used for immunoblotting assays with anti-GFP antibody. Degradation of the MdCIbHLH1 protein was accelerated in the presence of nitrate (Fig. 7A). In contrast, the MdCIbHLH1 proteins gradually accumulated in the absence of nitrate (Figure 7A). These results suggest that nitrate facilitates the degradation of MdCIbHLH1.

**Figure 7.**
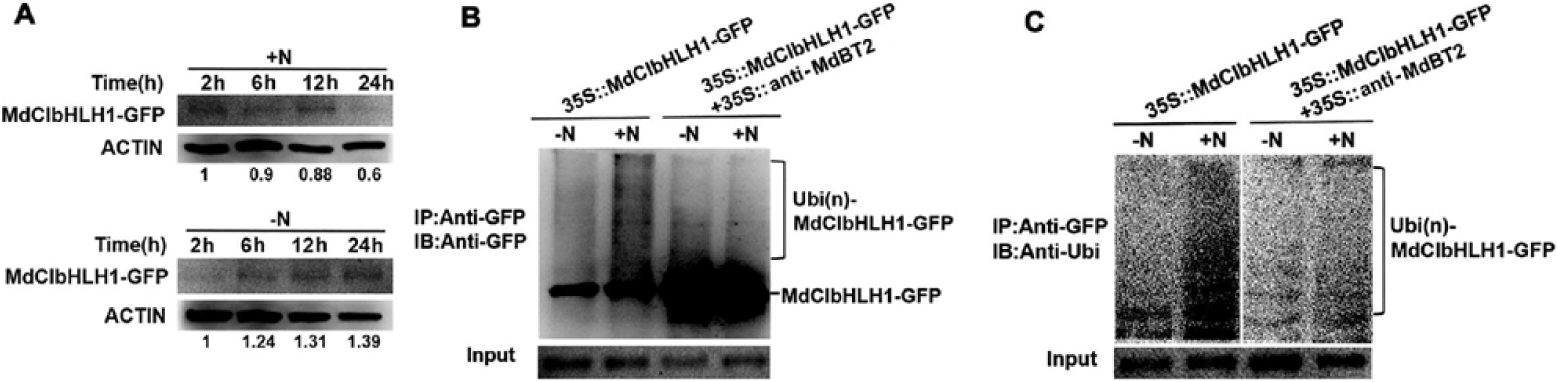
Nitrate participates in the regulation of MdBT2-mediated MdCIbHLH1 protein stability. **A,** The protein stability of MdCIbHLH1 in response to nitrate. 35S:: MdCIbHLH1-GFP were incubated without nitrate (-N) and with nitrate (+N) respectively. The total proteins extracted from these calli samples were then used for immunoblotting assays with anti-GFP antibody. **B and C,** The immunoblotting assays were conducted using 35S::MdCIbHLH1-GFP treated with or without nitrate.

To examine the effect of nitrate on MdCIbHLH1 protein ubiquitination, immunoblotting assays were conducted using 35S::MdCIbHLH1-GFP treated with or without nitrate. Exogenous nitrate treatment accelerated the ubiquitination of MdCIbHLH1 in 35S::MdCIbHLH1-GFP, while the MdCIbHLH1 protein ubiquitination was alleviated in the 35S::MdCIbHLH1-GFP + 35S::anti-MdBT2 (Fig. 7B). These results suggest that nitrate is involved in the regulation of MdBT2-mediated MdCIbHLH1 protein ubiquitination.

In addition, the level of the ubiquitinated MdCIbHLH1 protein was also assessed using an anti-ubi antibody that recognizes only ubiquitinated MdCIbHLH1 protein. Similarly, the ubiquitin signal was significantly enhanced in the 35S::MdCIbHLH1-GFP treated with nitrate compared with absence of nitrate, whereas the ubiquitin signal was reduced in the 35S::MdCIbHLH1-GFP + 35S::anti-MdBT2 (Fig. 7C)

Overall, these results suggest that nitrate acting as a signal regulates MdBT2-mediated MdCIbHLH1 protein stability.

### MdBT2 modulates malate accumulation in an MdCIbHLH1-dependent manner

To investigate whether MdBT2 and MdCIbHLH1 regulate malate accumulation in apple, a viral vector-based method was applied to alter their expression using vector TRV for suppression. Two viral constructs, including MdBT2-TRV and MdCIbHLH1-TRV, were obtained. These constructs including MdBT2-TRV and MdCIbHLH1-TRV, as well as two combinations MdBT2-TRV + MdCIbHLH1-TRV were used for fruit infiltration, with the empty vectors as controls (Fig. 8A). qPCR assays showed that the transcript levels of *MdBT2* and *MdCIbHLH1* genes were significantly decreased after being infiltrated with MdBT2-TRV and MdCIbHLH1-TRV (Fig. 8B).

**Figure 8.**
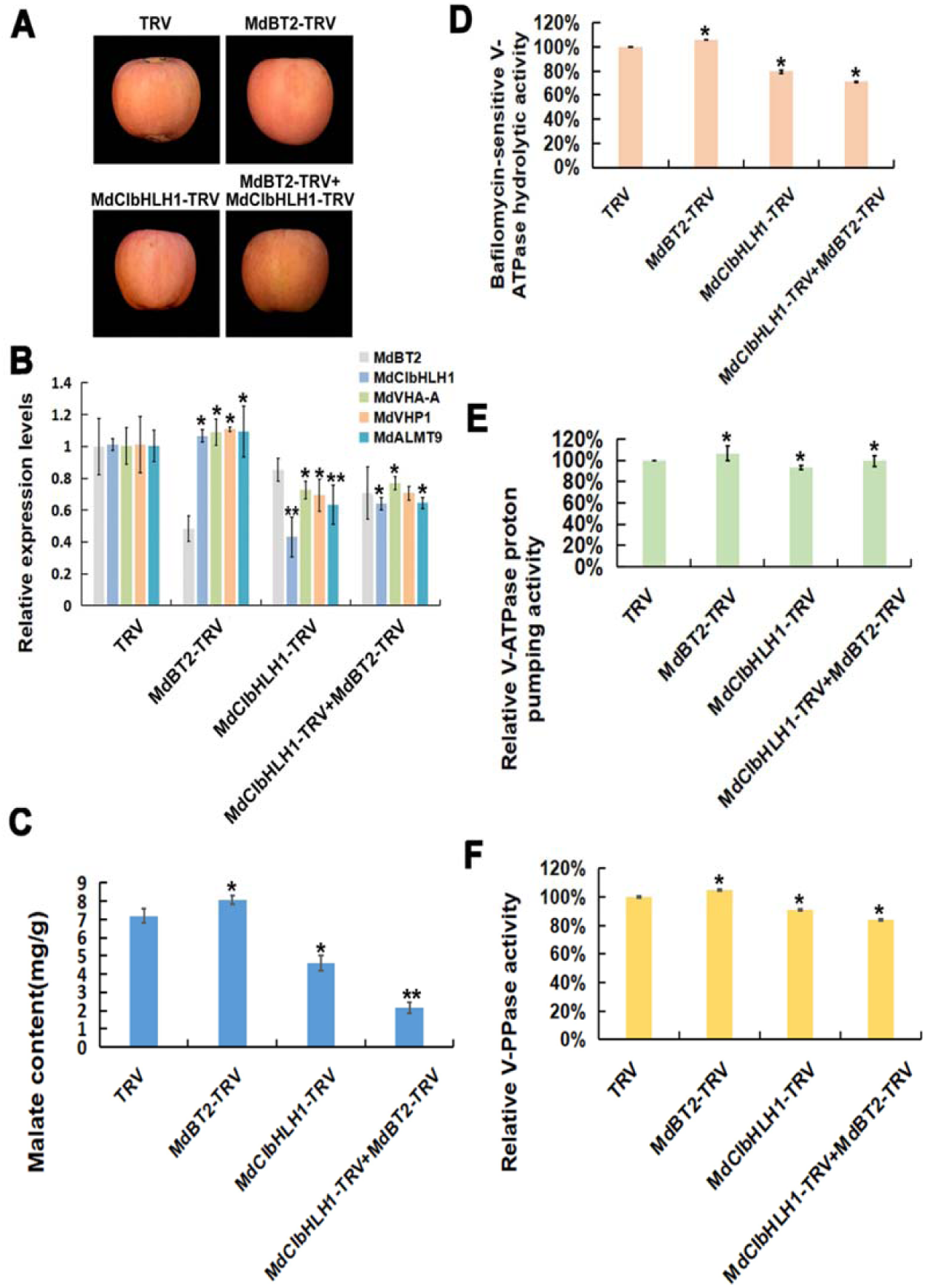
MdBT2 modulates malate accumulation in an MdCIbHLH1-dependent manner. **A,** Viral vector-based method was applied to apple injection assays. TRV, TRV1 + TRV2; MdBT2-TRV, TRV1 + MdBT2-TRV2; MdCIbHLH1-TRV, TRV1 + MdCIbHLH1-TRV2; MdBT2-TRV2 + MdCIbHLH1-TRV2, TRV1 + MdBT2-TRV2 + MdCIbHLH1-TRV2. **B,** The qPCR analysis of the expression of *MdVHA-A*, *MdVHP1* and *MdALMT9*. **C to F,** The malate contents, hydrolytic and proton-pumping activities of V-ATPase and V-PPase were measured in apple fruit tissues around the sites infiltrated with the different viral constructs. Significant difference was detected by *t*-test. *P < 0.05, **P < 0.01.

In response to suppression of *MdCIbHLH1*, transcript levels of the MdCIbHLH1-targeted genes involved in malate accumulation including *MdVHA-A*, *MdVHP1* and *MdALMT9*, were decreased. By contrast, the transcripts of these genes were increased in the MdBT2-TRV injected apple fruits (Fig. 8B). Subsequently, malate levels, hydrolytic and proton-pumping activities of V-ATPase and V-PPase activity were measured in apple fruit tissues around the infiltration sites. MdCIbHLH1-TRV injected apple fruits had lower malate levels and lower proton-pumping activities than the TRV control, whereas MdBT2-TRV injected apple fruits produced higher malate levels and proton-pumping activities than the control (Fig. 8, C-F). In the MdBT2-TRV + MdCIbHLH1-TRV double-injected apple fruits, *MdCIbHLH1* suppression abolished the effect of MdBT2-TRV infiltration on malate accumulation and proton-pumping activities, indicating that MdBT2 controls malate accumulation at least partially, if not entirely, via MdCIbHLH1 in apple. Similar results were obtained in *MdBT2* and *MdCIbHLH1* transgenic apple plantlets (Supplemental Fig. S3).

In earlier work, we showed that MdCIbHLH1 interacts with and activates MdMYB73 to modulate malate accumulation and vacuolar acidification by directly activating *MdALMT9* in apples (Hu et al., 2017). Because MdBT2 mediates the ubiquitination and degradation of the MdCIbHLH1 protein, it is reasonable to predict that MdBT2 influences the transcription level of MdMYB73-downstream gene *MdALMT9*. To verify this hypothesis, GUS assays were conducted to determine the effect of MdBT2 on the transcriptional activity of *MdALMT9*. (Supplemental Fig. S4). MdBT2-MdCIbHLH1 regulatory module negatively regulates GUS transcription of MdMYB73-downstream gene *MdALMT9*, which is driven by the *MdALMT9* promoter.

Taken together, these results further demonstrate that MdBT2 modulates malate accumulation in an MdCIbHLH1-dependent manner.

## Discussion

Malate is a crucial metabolite regulated by various signaling pathways that integrate responses of gene expression to environmental factors such as nutrient availability. Physiological studies have demonstrated that malate and nitrate have intricate regulation mechanisms during plant growth and development (Stitt, 1999; Etienne et al., 2013). Although a connection between nitrate and malate has long been speculated, the exact molecular mechanism has remained unclear. Some reports suggest that nitrogen positively regulates fruit acidity (Reitz and Koo, 1960; Ruhl, 1989; Jia et al., 1999; Radi et al., 2003), while others have shown a negative relationship between nitrate level and fruit acidity (Spironello et al., 2004; Wang et al., 2010). The present study reveals that high nitrate inhibits malate accumulation in apple (Fig. 1; Supplemental Fig. S1). MdBT2, a nitrate-responsive protein, which was shown to be involved in regulating anthocyanin accumulation through interacting with the MdMYB1 in our earlier work (Wang et al., 2018), plays a key role in modulating malate accumulation in response to nitrate. MdBT2 interacts with and ubiquitinates MdCIbHLH1 in response to nitrate (Fig. 2-8), providing the molecular link between nitrate availability and malate accumulation.

BT2 is a scaffold protein that recruits CUL3 protein to form the E3 ligase complex or bridges an unknown ubiquitin E3 ligase to ubiquitinate and degrade the target proteins (Robert et al., 2009; Zhao et al., 2016; Wang et al., 2018). MdCUL3 protein is required for the MdBT2-mediated ubiquitination and degradation of MdbHLH104 for iron homeostasis in apple, with the TAZ domain of MdBT2 being essential for the interaction (Zhao et al., 2016). However, MdBT2 promotes the ubiquitination and degradation of MdMYB1 in regulating anthocyanin biosynthesis through an MdCUL3-independent pathway, while the BACK domain of MdBT2 is indispensable for the interaction (Wang et al., 2018). In this study, both BTB domain and BACK domain are required for the interaction between MdBT2 and MdCIbHLH1 (Fig. 3). Based on these findings, we speculate that MdBT2 possibly ubiquitinates MdCIbHLH1 through an MdCUL3-independent pathway.

In apple, MdCIbHLH1, an ICE1-like protein, regulates cold tolerance in a CBF-dependent way (Feng et al., 2012). The ICE1-CBF-COR cold response pathway is one of the dominating cold signaling modules that bring about the cold tolerance response in *Arabidopsis* (Chinnusamy et al., 2003; Shi et al., 2018). It has been known that malate accumulation is affected by temperature. High temperature reduces malate accumulation in fruit (Buttrose et al., 1971; Lobit et al., 2006; Etienne et al., 2013) whereas low temperature facilitates vacuolar malate accumulation probably by affecting the proton pumps transport activity and membrane fluidity (Murata and Los, 1997; Terrier et al., 2001; Etienne et al., 2013). As malate serves as an osmoticum for plant resistance to low temperature (Ruffner et al., 1976; Friemert et al., 1988; Kovermann et al., 2007) and salinity (Hu et al., 2016b), our previous work (Hu et al., 2017) and current work strongly suggest that MdCIbHLH1 regulates plant cold tolerance via more than one pathway. In addition to the CBF-dependent pathway, MdCIbHLH1 controls cold tolerance via regulating malate accumulation by the MdCIbHLH1-MdMYB73 pathway, which was previously described in our study (Hu et al., 2017).

Malate accumulation involves transcriptional regulation of vacuolar proton pumps and malate transporters (Xie et al., 2012; Li et al., 2017; Hu et al., 2017). It has been reported that MdbHLH3 plays a central role in the accumulation of both malate and anthocyanins by interacting with MdMYB1 and binding to the promoter of MdMYB1 (Xie et al., 2012; Hu et al., 2016a). MdCIbHLH1 interacts with MdMYB73 and enhances its transcriptional activity on the downstream target genes in regulating malate accumulation (Fig. 4 and 8; Supplemental Fig. S4; Hu et al., 2017). In both cases, we speculate that the MBW complex is recruited to regulate malate accumulation. However, the target proteins of MdbHLH3 and MdCIbHLH1 differ in their functions in regulating malate vs. anthocyanin accumulation. Recent studies have identified *Noemi*, encoding a bHLH transcription factor, controls two co-evolved traits in citrus, fruit acidity and anthocyanin levels (Butelli et al., 2019; Zhu et al., 2019). Our findings here suggest that *Noemi* might regulate fruit acidity and anthocyanin levels via a pathway similar to MdbHLH3-MdMYB1 (Xie et al., 2012; Hu et al., 2016a).

Based on the findings presented in this work and elsewhere (Hu et al., 2017), we propose a working model for the regulation of malate accumulation by nitrate (Fig. 9). In this model, when not ubiquitinated under nitrate deficiency, MdCIbHLH1 enhances the activity of MdMYB73, which maintains the transcription of malate-associated genes at a high level, resulting in high malate accumulation. However, MdBT2 ubiquitinates and degrades MdCIbHLH1 in response to high nitrate, which reduces the transcription of malate-associated genes by decreasing the activity of MYB73, leading to lower malate accumulation. Our findings provide new insights into the molecular mechanism by which nitrate affects malate accumulation and fruit quality, providing the link between two key metabolites in plant metabolism. These findings not only help us understand the complex regulatory network of malate accumulation but also are of great significance in guiding molecular breeding programs as well as traditional apple breeding programs for fruit quality improvement.

**Figure 9.**
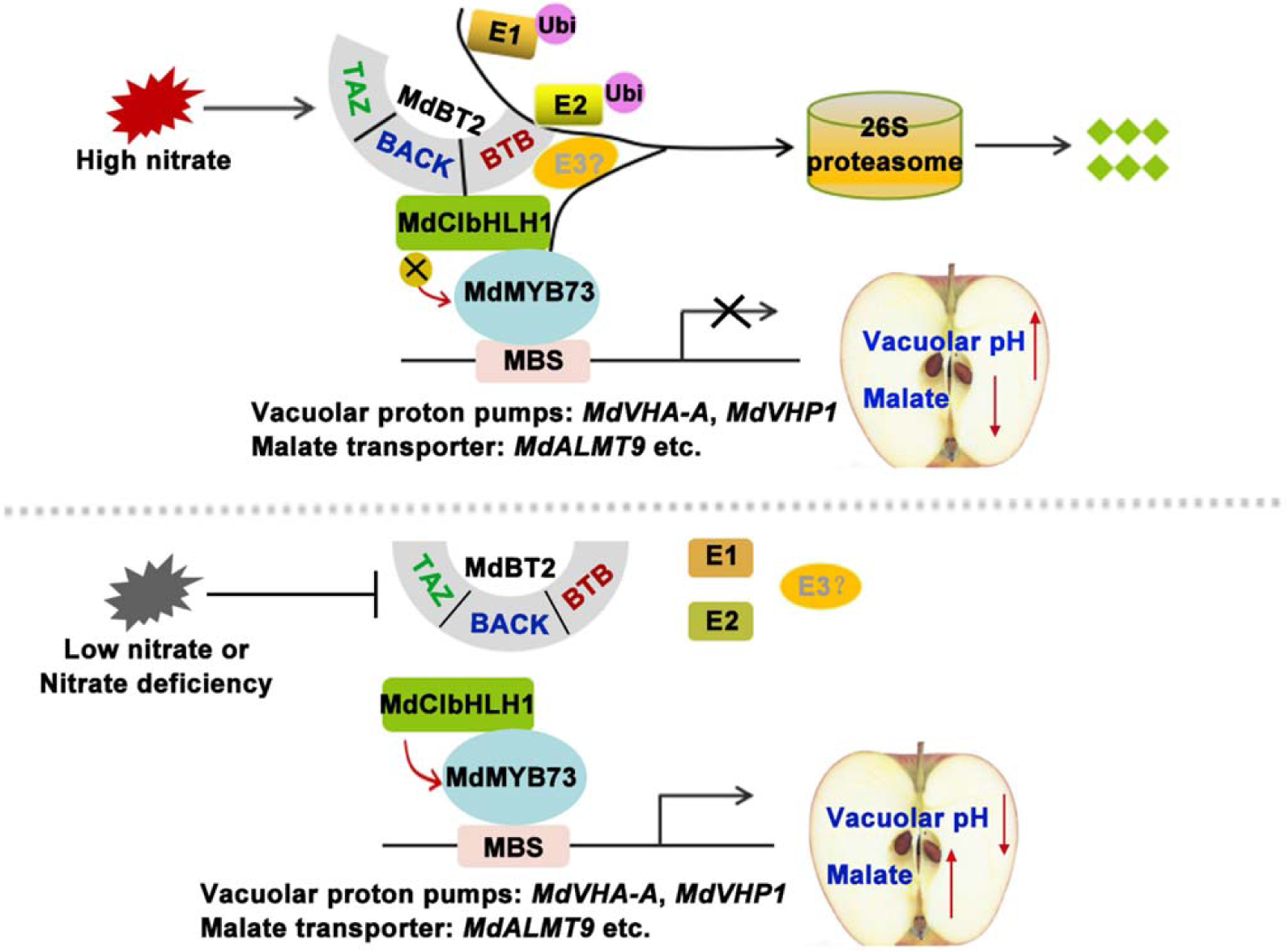
Work model demonstrating that BTB-TAZ protein MdBT2 ubiquitinates and degrades MdCIbHLH1 in response to high nitrate, whereas MdBT2 cannot ubiquitinate MdCIbHLH1 in low nitrate to regulate target gene activity of *MdMYB73* to modulate malate accumulation for fruit quality.

## Materials and methods

### Plant materials and treatments

‘*GL-3*’ apple (*Malus* × *domestica*) plantlets, ‘*Royal Gala*’ apple fruits and ‘*Orin*’ calli were used. The ‘*GL3*’ and transgenic apple plantlets were grown on Murashige-Skoog (MS) medium containing 0.2 mg L^-1^ NAA, 0.2 mg L^-1^ GA and 0.6 mg L^-1^ 6-BA at 25°C under long-day conditions (16 h light / 8 h dark). ‘*Orin*’ apple calli were grown on MS medium with 1.5 mg L^-1^ 2,4-D, and 0.4 mg L^-1^ 6-BA at 25°C in the dark. For phenotypic analyses of the nitrate-deficiency response, the ‘*GL-3*’ cultures and ‘*Orin*’ apple calli were transferred from nitrate-sufficient medium (MS medium with the indicated amount of KNO_3_, 5 mM) to nitrate-deficient medium (MS medium without nitrogen, followed by the indicated amount of KCl, 5 mM, as control), and then grown for 3 days.

### Measurement of malate contents

Samples were extracted with 95% (v/v) ice-cold methanol first and then 80% (v/v) ice-cold methanol after being frozen and ground in liquid nitrogen. The extracts were evaporated to dryness, dissolved in deionized water, and then centrifuged at 4000g. The supernatants were filtered through a 0.45 μm membrane filter and then analyzed with high performance liquid chromatography (HPLC) as described by Hu et al. (2016a).

### Activity assays of V-ATPase and V-PPase

Isolation of tonoplast membranes was performed as described by Terrier et al. (2001). The bafilomycin A_1_-sensitive ATP hydrolytic activity and V-ATPase H^+^ transport activity as well as V-PPase activity were measured as described by Hu et al. (2016a, 2017).

### Measurement of vacuolar pH

Isolation of protoplasts from apple fruit, plantlets and calli as well as vacuolar pH imaging were conducted as previously described by Hu et al. (2016a, 2017). A pH-sensitive fluorescent dye BCECF-AM was used to measure vacuolar pH. The vacuolar pH was quantified by the ratio of 488 nm and 458 nm excitation wavelengths. The ion concentration tool of Zeiss LSM confocal software was used to generate the ratio images.

### Quantitive RT-PCR (qPCR) assays

Total RNA was extracted from apple flesh, plantlets and calli using RNA Plant Plus Reagent (Tiangen, Beijing, China) and qPCR analysis was performed as described previously (Hu et al., 2019). The primers used for qPCR assays are listed in Supporting Information Table S1. The sequences used in this study were obtained from the GDR databases (https://www.rosaceae.org/). The GenBank accession numbers are: *MdCIbHLH1* (MDP0000662999), *MdBT2* (MDP0000151000), *MdMYB73* (MDP0000894463), *MdVHA-A* (MDP0000844729), *MdVHP1* (MDP0000688191), *MdALMT9* (MDP0000290997).

### Yeast two-hybrid (Y2H) assay

Y2H assays were performed as described by Wang et al. (2018). The full-length cDNA of MdCIbHLH1 was amplified and inserted into pGAD424, while MdBT2 cDNAs and truncated sequences (amino acids 1-335, 1-195, 1-128, 129-195, 129-335) were amplified and inserted into pGBT9. The plasmids of pGAD424-MdCIbHLH1 and the pGBT9-MdBT2s were co-transformed into Y2H Gold (Clontech, Mountain View, CA, USA). The yeasts were grown on -L/-T selection medium for the transformation control and on -L/-T/-H/-A selection medium with or without X-gal for the interaction analysis.

### Pull-down assays

The full-length cDNA of MdCIbHLH1 was amplified by PCR and inserted into pET-32a to generate HIS-tagged recombinant protein (MdCIbHLH1-HIS). The full-length cDNA of MdBT2 was also amplified by RT-PCR and cloned into pGEX-4T to produce GST-tagged fusion protein (MdBT2-GST). The fusion proteins of MdCIbHLH1-HIS, MdBT2-GST and empty GST were used for the pull-down assays according to Hu *et al.* (2019).

### Bimolecular fluorescence complementation (BiFC) assays

The full-length cDNA of MdBT2 was inserted into the vector 35S::pSPYNE-nYFP (MdBT2-nYFP) and MdCIbHLH1 was inserted into the vector 35S::pSPYCE-cYFP (MdCIbHLH1-cYFP). Solutions of *Agrobacterium* harboring recombinant plasmids (MdBT2-nYFP + MdCIbHLH1-cYFP) were injected into tobacco leaves. The empty vectors (MdBT2-nYFP + cYFP and nYFP + MdCIbHLH1-cYFP) were used as negative controls.

### Protein degradation and ubiquitination assays

For the degradation assays of the MdCIbHLH1 protein *in vitro*, apple calli (35S::MdBT2, WL and 35S::anti-MdBT2) and the MdCIbHLH1-HIS fusion protein were prepared. The extraction buffer contained 25 mM Tris (pH 7.5), 5 mM DTT, 10 mM NaCl, 10 mM MgCl_2_, 4 mM PMSF and 10 mM ATP. The MdCIbHLH1-HIS fusion protein and apple calli were extracted and incubated at 22°C. Samples were collected at the indicated time (0 h, 1 h, 2 h, 4 h and 6 h) and examined using an anti-HIS antibody. For the proteasome inhibitor experiments, 50 µM MG132 was added. For the degradation assays of the MdCIbHLH1 protein *in vivo*, two types of apple calli (35S::MdCIbHLH1-GFP and 35S::MdCIbHLH1-GFP + 35S::MdBT2-MYC) were prepared. Total proteins were monitored at the indicated times (0 h, 1 h, 2 h, 4 h and 6 h) when 250 mM CHX was added, and then the anti-GFP antibody was used for immunoblotting assays.

For the ubiquitination assays *in vivo*, three types of apple calli (35S::MdCIbHLH1-GFP, 35S::MdCIbHLH1-GFP + 35S::MdBT2-MYC and 35S::MdCIbHLH1-GFP + 35S::anti-MdBT2) were prepared and treated with 50 µM MG132 for 10 h before extraction. The protein extracts were immunoprecipitated using a Pierce Classic Protein A IP Kit (ThermoFisher Scientific, Waltham, MA, USA) with anti-GFP and anti-ubi antibodies. For the ubiquitination assays *in vitro*, 35S::MdBT2-MYC apple calli were treated with 50 µM MG132 for 10 h to obtain the MdBT2-MYC active protein using Pierce Classic Protein A IP Kit. The incubation buffer contained 50 mM Tris (pH 7.5), 2 mM DTT, 50 mM MgCl_2_, 2 mM ATP, 100 ng rabbit E1, 100 ng human E2 and 1 ug ubi. The buffer and MdCIbHLH1-HIS protein with or without MdBT2-MYC active protein *in vivo* were co-incubated at 30°C for 24 h. Protein ubiquitination was detected using anti-HIS and anti-ubi antibodies.

### Apple injection assays

Fruit injection assays were carried out as described previously (Hu et al., 2016). The CDSs of MdBT2 and MdCIbHLH1 were amplified and inserted into viral vector to obtain MdBT2-TRV (TRV1 + MdBT2-TRV2), MdCIbHLH1-TRV (TRV1 + MdCIbHLH1-TRV2) and MdBT2-TRV + MdCIbHLH1-TRV (TRV1 + MdBT2-TRV2 + MdCIbHLH1-TRV2), with the TRV vector (TRV1 + TRV2) used as control. The mixture of vectors and the *A. tumefaciens* solutions were injected into apple fruit peel and flesh and tissues around the injection hole.

### Statistical analysis

All experiments were performed in triplicate. Error bars show the standard deviation of three biological replicates. Significant difference was detected by *t*-test using G_RAPH_P_AD_ P_RISM_ 6.02 software (*, P < 0.05; **, P < 0.01).

## Supplemental Data

**Supplemental Fig. S1.** Malate contents under a series of nitrate concentrations in apple.

**Supplemental Fig. S2.** BTB/TAZ protein MdBT2 controls malate accumulation in response to nitrate.

**Supplemental Fig. S3.** MdBT2 modulates malate accumulation in an MdCIbHLH1-dependent manner in response to nitrate.

**Supplemental Fig. S4.** The MdBT2-MdCIbHLH1 regulatory module negatively regulates MdMYB73-downstream gene *MdALMT9*.

**Supplemental Table S1.** The primers used in this study.

## Acknowledgements

We would like to thank Prof. Takaya Moriguchi of National Institute of Fruit Tree Science, Japan, for ‘*Orin*’ apple calli. This work was supported by grants from the National Key Research and Development Program of China (2018YFD1000200), the National Natural Science Foundation of China (31972375, 31772288), the Ministry of Agriculture of China (CARS-28), Shandong Province (SDAIT-06-03), and Nanjing Agricultural University (ZW201805).

## Author Contributions

Y.-J.H. and D.-G.H. planned and designed the research. Q.-Y.Z., K.-D.G., J.-H.W., J.-Q.Y., X.-F.W. and C.-X.Y. performed experiments, conducted fieldwork, analysed data etc. Q.-Y.Z., D.-G.H., Y.-J.H. and L.C wrote the manuscript.

